# Four new species of *Cichlidogyrus* (Platyhelminthes, Monogenea, Dactylogyridae) from Lake Victoria haplochromine cichlid fishes, with the redescription of *C. bifurcatus* and *C. longipenis*

**DOI:** 10.1101/2021.01.29.428376

**Authors:** Tiziana P Gobbin, Maarten PM Vanhove, Ole Seehausen, Martine E Maan, Antoine Pariselle

**Author notes:** first two authors contributed equally (Tiziana P Gobbin, Maarten PM Vanhove).

## Abstract

African cichlids are model systems for evolutionary studies and for host-parasite interactions, because of their adaptive radiations and because they harbour many species of monogenean parasites with high host-specificity. Here, we sampled five locations in southern Lake Victoria, the youngest of the African Great Lakes. We surveyed gill-infecting monogeneans from 18 cichlid species belonging to the Lake Victoria radiation superflock and two cichlid species representing two older and distantly related lineages. We found one species of *Gyrodactylus* (Gyrodactylidae, Monogenea), *Gyrodactylus sturmbaueri* Vanhove, Snoeks, Volckaert & Huyse, 2011, and seven species of *Cichlidogyrus* (Dactylogyridae, Monogenea). Four species are herein described: *Cichlidogyrus pseudodossoui* n. sp., *C. nyanza* n. sp., *C. furu* n. sp., *C. vetusmolendarius* n. sp.. Another species is reported but not formally described (because of few specimens and morphological similarity with *C. furu* n. sp.). Two other species are redescribed: *Cichlidogyrus bifurcatus* Paperna, 1960 and *C. longipenis* Paperna & Thurston, 1969. Our results confirm that the monogenean fauna of Victorian littoral cichlids displays lower species richness and lower host-specificity than that of Lake Tanganyika littoral cichlids. In *C. furu* n. sp., hooks V are clearly longer than the other hooks, highlighting the need to re-evaluate the current classification system of haptoral configurations that considers hook pairs III-VII as rather uniform. Some morphological features of *C. bifurcatus, C. longipenis* and *C. nyanza* n. sp. suggest that these are closely related to other congeners that infect haplochromines. We also found morphological indications that representatives of *Cichlidogyrus* colonised Lake Victoria haplochromines or their ancestors at least twice, which is in line with the Lake Victoria superflock being colonized by two cichlid tribes (Haplochromini and Oreochromini).

**Disclaimer:** This preprint is disclaimed for purposes of Zoological Nomenclature in accordance with the International Code of Zoological Nomenclature, Fourth Edition Articles 8.2 and 8.3 (ICZN 1999). **No new names or nomenclatural changes are available from statements in this preprint.**

**Résumé - Quatre espèces nouvelles de *Cichlidogyrus* (Platyhelminthes, Monogenea, Dactylogyridae) parasites d’haplochrominé (Cichlidae) du lac Victoria, avec la redescription de *C. bifurcatus* and *C. longipenis*:** A cause des radiations adaptatives qu’ils ont subies, les cichlidés africain sont des systèmes modèles pour étudier l’évolution, mais aussi les relations hôtes/parasites, car ils hébergent de nombreuses espèces de Monogènes parasites qui présentent une spécificité étroite vis-à-vis de leurs hôtes. Dans ce travail, nous avons échantillonné cinq localités dans le Sud du lac Victoria, le plus jeune des grands lacs d’Afrique de l’Est. Nous avons examiné les Monogènes présents sur les branchies de 18 espèces de Cichlidés appartenant à la radiation adaptative « superflock » du lac Victoria et de deux espèces représentant deux lignées anciennes et non étroitement apparentées. Nous avons trouvé une espèce de *Gyrodactylus* (Gyrodactylidae, Monogenea), *Gyrodactylus sturmbaueri* Vanhove, Snoeks, Volckaert & Huyse, 2011 et sept espèces de *Cichlidogyrus* (Dactylogyridae, Monogenea). Quatre espèces nouvelles sont décrites dans le présent travail : *Cichlidogyrus pseudodossoui* n. sp., *C. nyanza* n. sp., *C. furu* n. sp., *C. vetusmolendarius* n. sp.. Une est signalée mais non décrite formellement (trop peux d’individus recueillis, morphologiquement proche de *C. furu* n. sp.). Deux autres sont redécrites : *Cichlidogyrus bifurcatus* Paperna, 1960 and *C. longipenis* Paperna & Thurston, 1969. Nos résultats confirment que la faune des Monogènes des Cichlidés du lac Victoria fait preuve d’une richesse spécifique et d’une spécificité moins importante que celle du lac Tanganyika. Chez *C. furu* n. sp. la paire de crochet V étant nettement plus longue que les autres, il faudra reconsidérer le système de classification actuel des types de hapteurs chez les *Cichlidogyrus,* qui considère que tous les crochets (III à VII) ont la même taille. Quelques caractéristiques morphologiques de *C. bifurcatus, C. longipenis* et *C. nyanza* n. sp. pourraient être la preuve d’une ascendance commune avec des congénères présents chez d’autres Haplochrominés. De même, certains caractères indiqueraient que des représentants des *Cichlidogyrus* ont colonisé les Haplochrominés du lac Victoria, ou leurs ancêtres, au moins à deux reprises, ce qui est cohérent avec une colonisation du lac par deux lignées de cichlidés distinctes (Haplochromini and Oreochromini).

## Introduction

Cichlid fish (Cichlidae) form one of the most species-rich families of vertebrates, occurring mainly in rivers and lakes in Africa and South America. They underwent spectacular adaptive radiations in many African lakes, including lakes Tanganyika, Malawi and Victoria [8, 20, 50]. The species flocks that evolved in the African Great Lakes display a large diversity in morphology, ecology and behaviour, and high levels of endemism [8, 54, 20, 60, 46, 61]. There are also many cases in which cichlids failed to radiate upon colonizing lakes [50, 60, 61]. Together, these characteristics have made African cichlids a rewarding model system for studying adaptation and speciation [21,20]. In recent years, evidence has accumulated that the diversity of cichlids in the African Great Lakes is also associated with a diversity of parasites [41, 57, 19, 12]. Monogenean flatworms are promising model parasites to study whether and how the diversification processes in cichlids and their parasites influence each other [38, 56]. This is because of their species richness in African cichlids, their narrow host-specificity compared to other cichlid parasites, and their direct lifecycle. Most studies of monogenean parasites of Great Lake cichlids have focused on Lake Tanganyika, the oldest of the three Great Lakes that also has by far the oldest cichlid radiations (e.g. [39, 57]). From this lake, 39 species of *Cichlidogyrus* Paperna, 1960 [32, 43] and three species of *Gyrodactylus* von Nordmann, 1832 [58] have been described from cichlids. Of these, only one species from each genus has been found also on non-Tanganyikan hosts: *Cichlidogyrus mbirizei* Muterezi Bukinga, Vanhove, Van Steenberge & Pariselle, 2012 and *Gyrodactylus sturmbaueri* Vanhove, Snoeks, Volckaert & Huyse, 2011. They were reported for the first time outside of Lake Tanganyika by Lerssutthichawal et al. [22] in a cultured *Oreochromis* hybrid in Thailand, and by Zahradníčková et al. [64] in *Pseudocrenilabrus philander* in Zimbabwe and South Africa, respectively. Conversely, only two monogenean species previously known from other cichlids outside Lake Tanganyika have been observed in a cichlid endemic to the Tanganyika basin: *Cichlidogyrus halli* (Price & Kirk, 1967) and *Scutogyrus longicornis* (Paperna & Thurston, 1969), both on *Oreochromis tanganicae* (Günther 1984). This indicates that the monogenean assemblage of Tanganyika cichlids is quite distinct from the parasite fauna in other cichlids. Here, we focus on Lake Victoria, where the monogenean fauna has been investigated less extensively. Research on its monogeneans peaked already in the 1960s-70s [34, 39], yielding ten species of *Cichlidogyrus,* two species of *Gyrodactylus* and a single species of *Scutogyrus,* all of which are found also on cichlids outside the Lake Victoria region, with the exception of *Cichlidogyrus longipenis* Paperna & Thurston, 1969. Only recently, Lake Victoria’s monogeneans received attention again, namely in ecological parasitology [24, 23, 19, 12, 13]. Two of these studies distinguished *Cichlidogyrus* at the species level [12, 13] and suggest that the species richness, level of endemism, and host-specificity of cichlid-infecting monogeneans may be lower in Lake Victoria than in Lake Tanganyika, a feature that Pariselle et al. [39] suggested to be linked to the younger age of the Lake Victoria cichlid species flock.

Lake Victoria is the youngest, and also the shallowest and most turbid of the three Great Lakes. It was completely dry until about 14’600 years ago [17]. Most of its current cichlid fauna evolved *in situ* after that dry period [16, 53, 62, 27]. The Lake Victoria cichlid superflock evolved from a hybrid swarm derived from at least two riverine lineages that colonized the lake [51, 27]. This provided the genetic variation that, together with ample ecological opportunities, allowed rapid speciation and adaptive radiation [49, 45, 25]. Cichlid species display a wide range of trophic specializations [15, 63, 48, 2, 3, 26, 44]. All Victorian haplochromines are female mouthbrooders [52] and many are rock-dwellers [48]. Older cichlid lineages (only distantly related to the species that radiated in the lake) also colonized the lake, but did not speciate: *Astatoreochromis alluaudi* Pellegrin, 1904, *Pseudocrenilabrus multicolor* (Schöller, 1903), *Oreochromis variabilis* (Boulenger, 1906) and *Oreochromis esculentus* (Graham, 1928). These are currently sympatric with species of the Lake Victoria superflock. Here, for the first time in over 40 years, we systematically survey the monogenean fauna infecting the three anciently divergent haplochromine cichlid lineages of Lake Victoria: the radiation lineage (represented by 18 species) and the two lineages that did not radiate *(Astatoreochromis alluaudi* and *Pseudocrenilabrus multicolor).* All the species sampled are host to *Cichlidogyrus* spp. (Monogenea), *Lamproglena monodi* (Copepoda), *Ergasilus lamellifer* (Copepoda), and glochidia larvae of freshwater mussels (Bivalvia) attached to their gills [19, 12]. Of the hosts, *A. alluaudi*has previously been investigated in taxonomic work for its monogenean gill parasites, next to nine members of the haplochromine radiation and four non-haplochromine cichlid species present in Lake Victoria (overview in [39]). Hence, our survey more than doubles the number of cichlid species from Lake Victoria that have been scrutinised and their monogenean parasites identified to species level or formally described. The present substantial expansion of host coverage, although not including a formal description at the time, allowed Gobbin et al. [12] to propose several patterns with regard to the diversity of *Cichlidogyrus* in Lake Victoria haplochromines. First, the non-radiating haplochromines harbour a monogenean fauna that is distinct from their radiating counterparts. This is not surprising, given that the non-radiating *A. alluaudi* is the sole host known for the only species of *Cichlidogyrus* that is currently known to be endemic to Lake Victoria, *C. longipenis.* Second, the dactylogyrid monogeneans infecting members of the haplochromine radiation were rarely found also on cichlids not belonging to the radiation, in line with the trends observed in the overview of Pariselle et al. [39]. The present study supports these observations with four formal taxonomic descriptions and two redescriptions.

## Materials and Methods

Fish were caught by angling and with gillnets of variable mesh sizes at five locations in southern Lake Victoria, Tanzania (**Table 1**, **Fig. 1**, **Table S.1**): the rocky islands Makobe (in May-August 2010 and June-October 2014), Kissenda, Python and Luanso and the swampy inlet stream Sweya (these four in June-October 2014). At Makobe (−2.3654, 32.9228), we collected 13 cichlid species belonging to the Lake Victoria radiation lineage and one species *(Astatoreochromis alluaudi)* representing an older lineage that did not speciate. At Sweya (−2.5841, 32.8970), we collected *A. alluaudi* as well as *Pseudocrenilabrus multicolor victoriae,* representing the other lineage that did not diversify. At Kissenda (−2.5494, 32.8276) and Python (−2.6237, 32.8567), we collected three additional species of the Lake Victoria radiation lineage. Two of those *(Pundamilia* sp. ‘pundamilia-like’ and *P.* sp. ‘nyererei-like’) are closely related to *P. pundamilia* and *P. nyererei* that occur at Makobe. At Luanso (−2.6889, 32.8842), we collected one species *(Pundamilia* sp. ‘Luanso’). These species pairs of *Pundamilia* represent replicate speciation events [28]. The sampled cichlid species vary in their micro-habitat and trophic specializations [15, 63, 48, 2, 3] and also in the extent of genetic differentiation [28, 29]. Within the radiation, divergence is 14’600 years old, while the divergence between both nonradiating lineages, and between them and the ancestors of the radiations in Lake Victoria, Lake Malawi and other lakes, is 8-10 million years old [28, 29, 44]. Fish were immediately sacrificed with an overdose of 2-phenoxyethanol, numbered and preserved in ethanol (some directly preserved in 100% ethanol, others fixed in 4% formalin and then transferred to 70% ethanol). In the laboratory, the right gill arches were removed and screened for macroparasites by inspecting gill filaments with a mounted needle, under a dissecting stereoscope (Zeiss Stemi 2000). Monogeneans were detached with tweezers and individually stored in 100% ethanol. Specimens were individually mounted onto a slide, treated with 20% sodium dodecyl sulphate (SDS) to soften tissues and then fixed in Hoyer’s medium. Specimens of *Cichlidogyrus* were examined with a phase-contrast microscope (Olympus BX41TF) and sclerotized parts (nomenclature and numbering according to ICOPA IV; [6]) were measured with Olympus Stream Essentials v. 1.9 software (**Fig. 2**, all measurements given in μm). Drawings of sclerotized parts were made with CorelDraw 2019 software on the basis of microphotographs taken with a Leica DM2500 microscope and LAS 6.0 software. Type and voucher specimens of the parasites were deposited in the Muséum national d’Histoire naturelle, Paris, France (MNHN), the Royal Museum for Central Africa, Tervuren, Belgium (RMCA) and the Iziko South African Museum, Cape Town, Republic of South Africa (SAMC); symbio(para)types and host vouchers (terminology: see [4]) are stored at the Swiss Federal Institute of Aquatic Science and Technology, Kastanienbaum, Switzerland (EAWAG). For more details on the host-parasite combinations and geographical range of the species of *Cichlidogyrus* used for differential diagnosis, see Scholz et al. [47] and Cruz-Laufer et al. [5].

**Figure 6.2.**
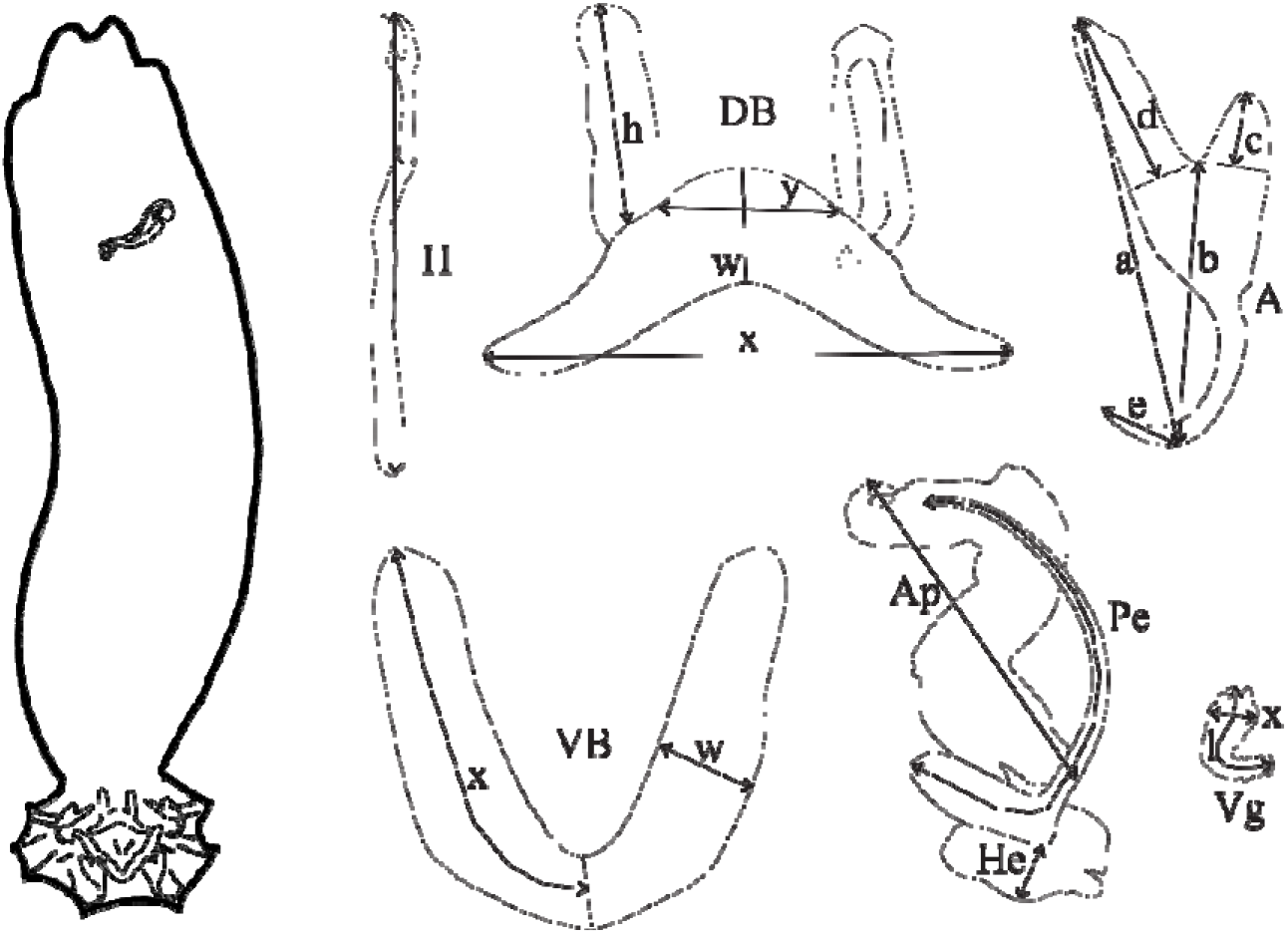
Schematic overview of a specimen of *Cichlidogyrus* with its typical morphological sclerotized structures (male copulatory complex in the upper part of the parasite image, attachment organ in the bottom part) and measurements used in the descriptions of the new species. Abbreviations: **A,** anchor **(a,** total length; **b,** blade length; **c,** shaft length; **d,** guard length; **e,** point length); **DB,** dorsal bar **(h,** auricle length; **w,** maximum straight width; x, total length; **y,** distance between auricles); **VB,** ventral bar (x, length of one ventral bar branch; **w,** maximum width); **H,** hook length; **Pe,** penis curved length; **He,** heel straight length; **Ap,** accessory piece straight length; **Vg,** vagina (**l**, vagina curved length; **x**, width).

**Table 1.** Haplochromine host species sampled in May-August 2010 and June-October 2014 at five localities (Makobe, Sweya, Kissenda, Luanso, Python) in southern Lake Victoria and the sample size of identified specimens of *Cichlidogyrus.* The radiation lineage is labelled with a circle (⍰), the two lineages that did not speciate are labelled with a square (∎) and a diamond (◆).

| Host sp. | Locality | n *Cichlidogyrus* | n fish |
| --- | --- | --- | --- |
| ■ *Astatoreochromis alluaudi* | Makobe | 106 | 8 |
| ■ *Astatoreochromis alluaudi* | Sweya | 19 | 3 |
| ◆ *Pseudocrenilabrus multicolor victoriae* | Sweya | 12 | 5 |
| ○ *"Haplochromis" cyaneus* | Makobe | 16 | 5 |
| ○ *Astatotilapia nubila* | Sweya | 6 | 1 |
| ○ *Labrochromis* sp. 'stone' | Makobe | 3 | 2 |
| ○ *Mbipia lutea* | Makobe | 14 | 3 |
| ○ *Mbipia mbipi* | Makobe | 26 | 11 |
| ○ *Neochromis gigas* | Makobe | 15 | 3 |
| ○ *Neochromis omnicaeruleus* | Makobe | 58 | 10 |
| ○ *Neochromis rufocaudalis* | Makobe | 13 | 4 |
| ○ *Neochromis* sp. 'unicuspid scraper' | Makobe | 46 | 18 |
| ○ *Paralabidochromis chilotes* | Makobe | 5 | 4 |
| ○ *Ptyochromis* sp. 'striped rock sheller' | Makobe | 3 | 1 |
| ○ *Pundamilia nyererei* | Makobe | 42 | 17 |
| ○ *Pundamilia pundamilia* | Makobe | 50 | 20 |
| ○ *Pundamilia* sp. 'pink anal' | Makobe | 21 | 6 |
| ○ *Ptyochromis xenognathus* | Kissenda | 18 | 4 |
| ○ *Pundamilia* sp. 'nyererei-like' | Kissenda | 29 | 11 |
| ○ *Pundamilia* sp. 'pundamilia-like' | Kissenda | 27 | 12 |
| ○ *Pundamilia* sp. 'Luanso' (blue form) | Luanso | 24 | 6 |
| ○ *Pundamilia* sp. 'Luanso' (red form) | Luanso | 28 | 7 |
| ○ *Pundamilia* sp. 'pundamilia-like' | Python | 2 | 1 |

This preprint is not a publication according to the ICZN, and especially according the emended Article 8 of the ICZN (ICZN 1999).

## Results

Pending a global revision of the species-rich genus *Cichlidogyrus* using genetic data (130 valid species described today: [37, 42, 43, 10]), the species belonging to this genus are grouped together by morphological similarities to facilitate their systematic analysis [40, 59]. Closely related species within the genus can be grouped by the morphology of their reproductive apparatus, whereas genera and subgeneric categories are more easily grouped by the morphology of their haptoral hard parts, one of the first criteria being the relative length of the hooks (see [36, 37]). During this survey, eight species of Monogenea were found on the gills of studied hosts. With the exception of a single specimen of *Gyrodactylus sturmbaueri* infecting *Ptyochromis xenognathus* from Kissenda (reported in [12] and deposited under accession number MRAC_VERMES_43410), all monogeneans corresponded to the diagnosis of *Cichlidogyrus* given in Pariselle & Euzet [37]. Two of the identified species of *Cichlidogyrus* were already known, the four others are herein formally described, and we characterise a potential fifth new species, for which we refrain from formal description for want of sufficient specimens.

### *Cichlidogyrus bifurcatus* Paperna, 1960

*Type host: Astatotilapia flaviijosephi* (Lortet, 1883).

*Type locality:* Sea of Galilee (spelled Sea of Gallilee in the original description).

*Site:* Gills.

*Additional hosts and localities:* [32]: young *Oreochromis aureus* (L.) from the Sea of Galilee (initially identified as *O. niloticus).* [34]: *Harpagochromis squamipinnis* Regan, 1921 and *“Haplochromis”* sp. ‘aeneocolor’ Greenwood, 1973 from Lake George, and Kazinga channel; “*H.” elegans* Trewavas, 1933 and *Haplochromis limax* Trewavas, 1933 from Lake George; *“H.”* sp. from Lake Edward and Entebbe, Lake Victoria; *Pseudocrenilabrus multicolor* (Schöller, 1903) from Lake Mulehe and a stream near Masindi, Lake Albert system, Uganda.

*New locality:* Lake Victoria, Sweya swampy inlet stream (−2.5841, 32.8970).

*New hosts: Pseudocrenilabrus multicolor victoriae* Seegers 1990; *Astatotilapia nubila* (Boulenger 1906).

*Infection parameters:* 3 of 6 *Pseudocrenilabrus multicolor victoriae* from Sweya infected with 2 individuals each, 1 of 1 *Astatotilapia nubila* from Sweya infected with 1 individual.

*Material:* Seven whole-mounted specimens in Hoyer’s solution.

*Voucher specimen:* MHNHxxxx-xx.

*Voucher host: Pseudocrenilabrus multicolor victoriae* Seegers 1990 from Sweya (EAWAG ID 109436, **Table S1**).

*Description* (**Table 2**, **Fig. 3**): Two pairs of anchors of equal size, with guard approximately 2 times as long as shaft. Dorsal anchors with guard and shaft more asymmetrical and blade shorter than ventral anchors. Ventral transverse bar V-shaped, with 2 branches with wing-shaped attachments along distal half. Dorsal transverse bar thin, tapering towards its extremities, and 2 small auricles inserted at its dorsal face. Hooks 7 pairs; I short (i.e. less than 1.7 times the length of II); III to VII on average short (i.e. less than 2 times the length of II) (see [36, 37]). Male copulatory complex (MCC) consisting of slightly curved penis with large basal bulb and constant diameter; accessory piece simple, ending in a fork with two smooth finger-like appendages of unequal size; developed heel with crenelated distal edge. Vagina not sclerotised.

**Figure 6.3.**
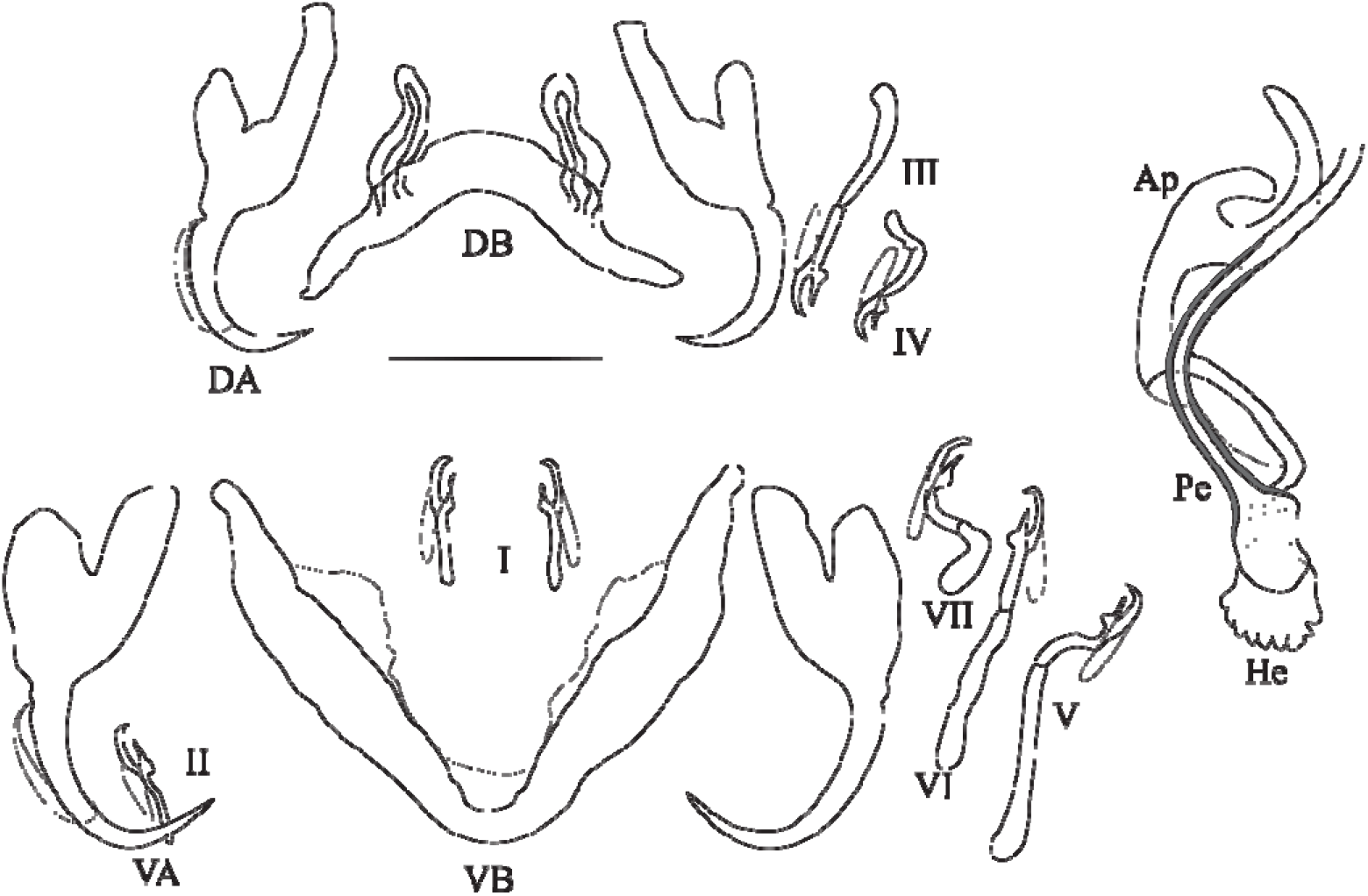
Sclerotized parts (haptor and male copulatory complex) of *Cichlidogyrus bifurcatus.* **I–VII,** hook pairs; **DA,** dorsal anchors; **DB,** dorsal transverse bar, **VA,** ventral anchors; **VB,** ventral transverse bar. Male copulatory complex: **AP,** accessory piece; **Pe,** penis;. Scale bar: 20 μm.

**Table 2.**
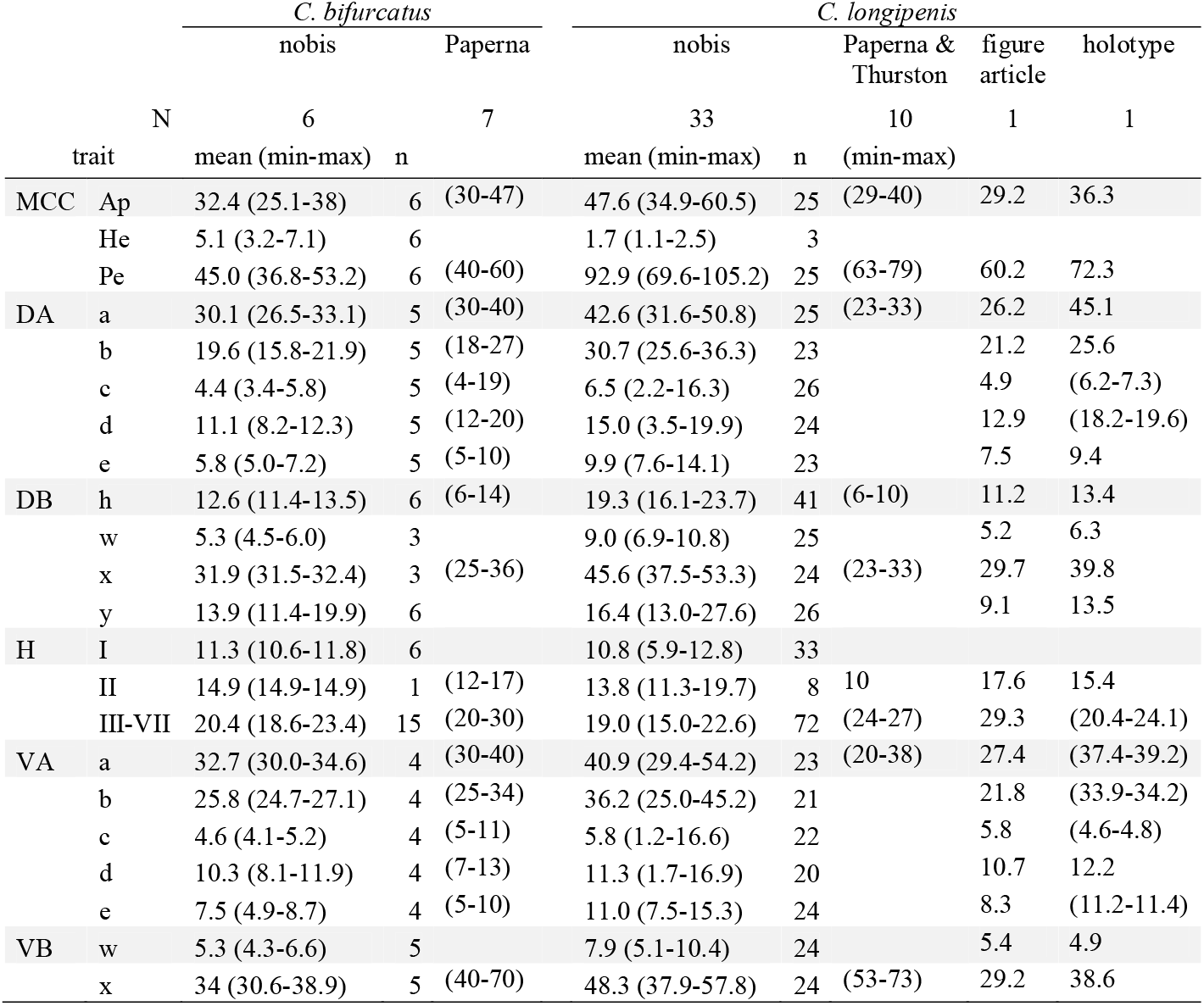
Measurements of *Cichlidogyrus bifurcatus* and *C. longipenis* redescribed from haplochromine host species of Lake Victoria (N = number of specimens, n = number of observations per trait per species) compared to those provided by Paperna [2] and Paperna & Thurston [3] (expressed as ranges only, because means were not reported in the original descriptions). For *C. longipenis,* we also report measurements extrapolated from the drawing of the original description and our measurements from the holotype. All measurements (terminology following ICOPA IV, [1]) in μm, given as the mean and range (in parentheses).

*Remarks:* the specimens found in Lake Victoria on *P. multicolor victoriae* resemble *C. bifurcatus* described by Paperna in 1960 on *A. flaviijosephi* in the Sea of Galilee by the shape and size of the MCC, that of the hooks, anchors and transverse bars. Only some minor differences in the size of some sclerotised parts are apparent (see **Table 2**), that may be due to two reasons, see remarks below under *C. longipenis. Cichlidogyrus bifurcatus* Paperna, 1960 was reported as *Cichlidogyrus* sp. VI in [12, 13].

### *Cichlidogyrus longipenis* Paperna & Thurston, 1969

*Type host: Astatoreochromis alluaudi* Pellegrin, 1904.

*Type locality:* Jinja, Uganda.

*Site:* Gills.

*New localities:* Lake Victoria, Makobe Island (−2.3654, 32.9228), Sweya swampy inlet (−2.5841, 32.8970).

*Hosts:* on type host and *Pundamilia* sp. ‘pink anal’.

*Infection parameters:* 9 of 9 *Astatoreochromis alluaudi* from Makobe infected with 2-17 individuals (mean intensity 10.7), 4 of 4 *A. alluaudi* from Sweya infected with 1-14 individuals (mean intensity 4.5), 2 of 7 *Pundamilia* sp. ‘pink anal’ from Makobe infected with 1 individual each.

*Material:* 33 whole-mounted specimens in Hoyer’s solution and the holotype MRAC M.T. 35.921.

*Voucher specimens:* MHNHxxxx-xx, RMCA_VERMES_43419, SAMC-A092084.

*Voucher hosts: Astatoreochromis alluaudi* Pellegrin, 1904 from Makobe (EAWAG ID 103148 and 103567, **Table S1**).

*Description* (**Table 2**, **Fig. 4**): Two pairs of anchors of equal size and unequal shape (guard more developed in dorsal anchors). Ventral transverse bar V-shaped. Dorsal transverse bar with two auricles inserted at its dorsal face. Hooks 7 pairs; I and III to VII short, except V of medium size (see [36, 37]). Penis long, wavy, thinner at proximal and distal extremities, rounded basal bulb, no heel; simple accessory piece, attached to the basal bulb of the penis by a filament, ending in a curved hook. No sclerotised vagina.

**Figure 6.4.**
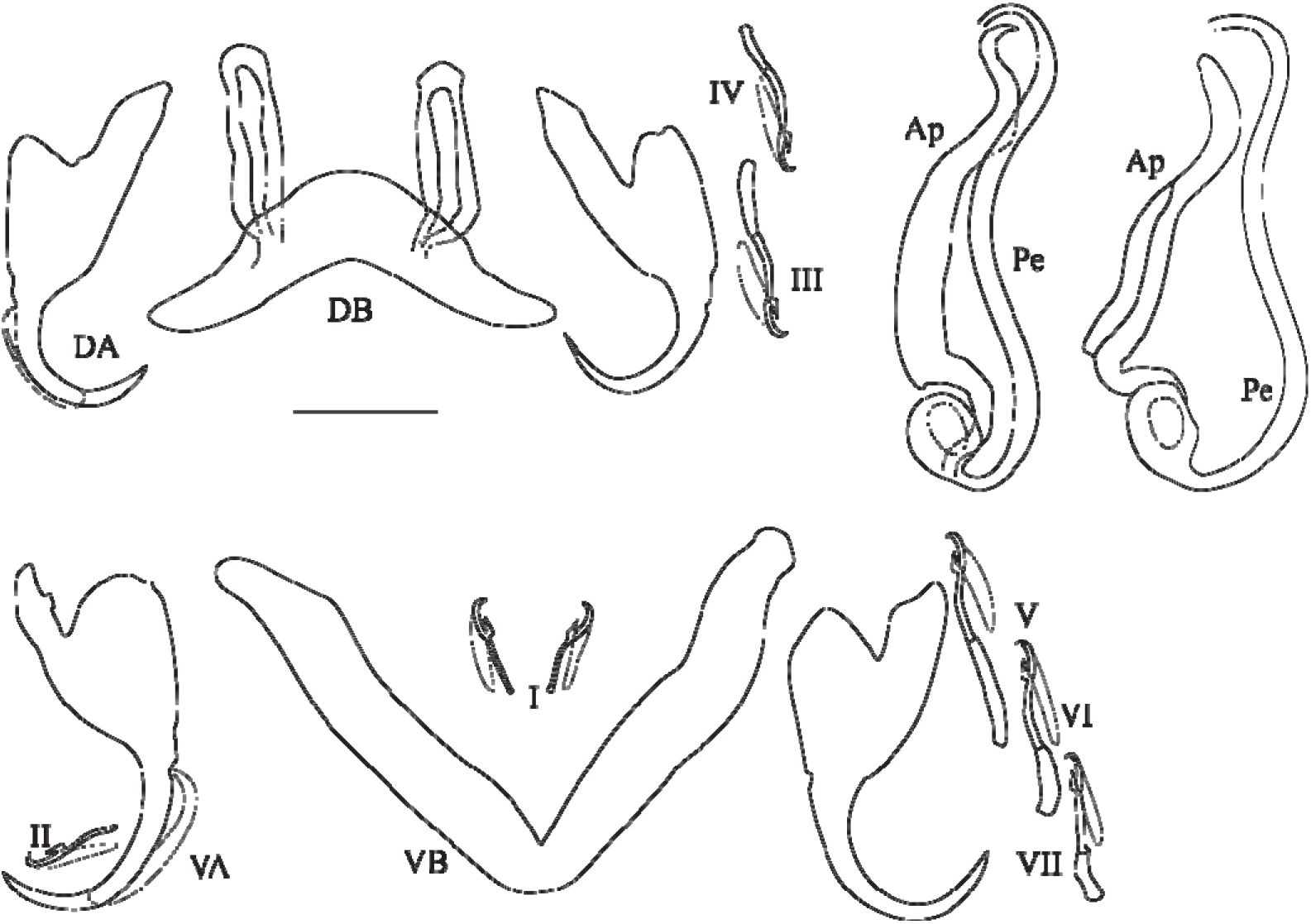
Sclerotized parts (haptor and male copulatory complex) of *Cichlidogyrus longipenis.* **I–VII**, hook pairs; **DA**, dorsal anchors; **DB**, dorsal transverse bar; **VA**, ventral anchors; **VB**, ventral transverse bar. Male copulatory complex (on the left as observed in most of our specimens, on the right the phenotype similar to the one reported by Paperna, 1960): **Ap**, accessory piece; **Pe**, penis. Scale bar: 20 μm.

*Remarks:* the specimens herein described share the combination of the following characters only with *C. longipenis* [35]: hooks pair I and II to VII short (or of medium size), penis long (>60 μm) with simple accessory piece and no heel, no sclerotised vagina. The measurements of some of these characters show differences between the original description and our data (**Table 2**). Some of our measurements were smaller when compared to those derived from the figure by Paperna & Thurston [35] (hook pairs II and III-IV) and to those reported in the original description (hook pairs II and III-IV, ventral bar length), but some of our measures were larger when compared to values reported in the original description, for example, the length of the accessory piece (29-40 vs 35-61, respectively), penis (63-79 vs. 70-105), ventral (20-38 vs. 29-54) and dorsal anchors (23-33 vs. 32-51). Even so, we are confident of our identification of these specimens as *C. longipenis:* as already quoted above, species delimitations in *Cichlidogyrus* are mainly based on the morphology of the reproductive apparatus [40, 59], which is similar in size and shape in our specimens compared to that of the type (**Fig. 4**, **Fig. 5**). The observed size differences may be due to two factors: 1) the mounting medium used when making the slides: Fankoua et al. [7] demonstrated that the use of Hoyer’s increases the size and modifies the shape of sclerotized parts, and/or 2) the way Paperna & Thurston [35] made the drawings and took the measurements for the original description. Although measurements given in the original description are compatible with its drawings and scale bars (**Table 2**), there are significant differences between these measurements and those we took from the type slide (**Table 2**) (an exception is the ventral bar total length, which seems incomparable: we measured the length of a branch while the original authors probably measured the total length of the bar). This is probably because Paperna & Thurston [35] did not use a camera lucida and made their drawings freehand, which may have led to magnification errors. *Cichlidogyrus longipenis* was reported as *Cichlidogyrus* sp. IV in [12, 13].

**Figure 6.5.**
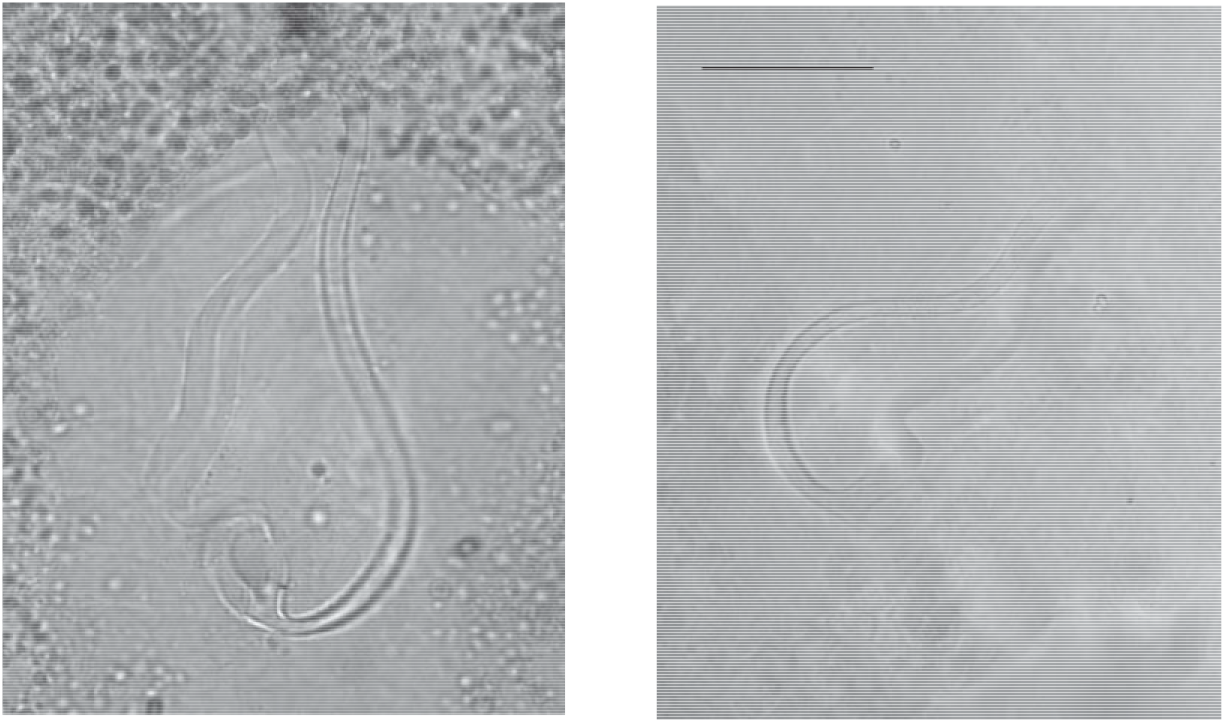
Micrographs of the male copulatory organ of *Cichlidogyrus longipenis,* from a parasite individual of the present study **(left)** and from the holotype by Paperna & Thurston,, 1969 **(right),** fixed in two different mediums. Scale bar: 20 μm.

### *Cichlidogyrus pseudodossoui* n. sp

urn:lsid:zoobank.org:act:2AA5987B-7AE1-407E-8E58-CB6CEFC89360

*Type host: Astatoreochromis alluaudi* Pellegrin, 1904.

*Other hosts: Ptyochromis xenognathus* (Greenwood, 1957); *Pundamilia pundamilia* Seehausen & Bouton, 1998; *P.* sp. ‘pundamilia-like’; *P. nyererei* (Witte-Maas & Witte, 1985); *P.* sp. ‘nyererei-like’; *Neochromis omnicaeruleus* Seehausen & Bouton, 1998; *N.* sp. ‘unicuspid scraper’.

*Infection parameters:* 5 of 9 *Astatoreochromis alluaudi* from Makobe infected with 1-2 individuals (mean intensity 1.2), 1 of 4 *A. alluaudi* from Sweya infected with 1 individual, 1 of 5 *Ptyochromis xenognathus* from Kissenda infected with 1 individual, 1 of 13 *Neochromis omnicaeruleus* from Makobe infected with 1 individual, 1 of 22 *N.* sp. ‘unicuspid scraper’ from Makobe infected with 1 individual, 1 of 23 *Pundamilia pundamilia* from Makobe infected with 1 individual, 1 of 14 *P.* sp. ‘pundamilia-like’ from Kissenda infected with 1 individual, 1 of 1 *P.* sp. ‘pundamilia-like’ from Python infected with 2 individuals, 3 of 24 *P. nyererei* from Makobe infected with 1-2 individuals (mean intensity 1.6), 1 of 16 *P.* sp. ‘nyererei-like’ from Kissenda infected with 2 individuals.

*Site:* Gills.

*Type locality:* Makobe Island (−2.3654, 32.9228).

*Other localities:* Kissenda Island (−2.5494, 32.8276), Python Island (−2.6237, 32.8567), Sweya swampy inlet (−2.5841, 32.8970).

*Material:* 10 whole-mounted specimens in Hoyer’s solution.

*Holotype:* MHNHxxxx-xx.

*Paratypes:* MHNHxxxx-xx, RMCA_VERMES_43421, SAMC-A092086.

*Symbiotype: Astatoreochromis alluaudi* Pellegrin, 1904 from Makobe (EAWAG ID 103148, **Table S1**).

*Symbioparatypes: Astatoreochromis alluaudi* Pellegrin, 1904 from Makobe (EAWAG ID 103571, **Table S1**); *Neochromis omnicaeruleus* Seehausen & Bouton, 1998 from Makobe (EAWAG ID 105655, **Table S1**); *Ptyochromis xenognathus* (Greenwood, 1957) from Kissenda (EAWAG ID 12306, **Table S1**).

*Etymology:* The species epithet refers to the similarity of this newly described species with its congener *C. dossoui* Douёllou 1993.

*Description* (**Table 3**, **Fig. 6**): Two pairs of anchors of unequal size (ventral larger) and similar shape: short blade, and shaft and guard symmetrical in appearance and of similar size. Ventral transverse bar V-shaped (rounded on its anterior edge). Dorsal transverse bar with two developed auricles inserted at its dorsal surface. Hooks 7 pairs; I short; III to VII long (see [36, 37]). MCC consisting of a J-shaped and thin penis starting at right angle from an ovoid bulb; accessory piece thick, Z-shaped, with a recurved tooth at its distal extremity; developed rounded heel. Vagina thick walled, bent at right angle, with annulated proximal third, broadened towards distal extremity.

**Figure 6.6.**
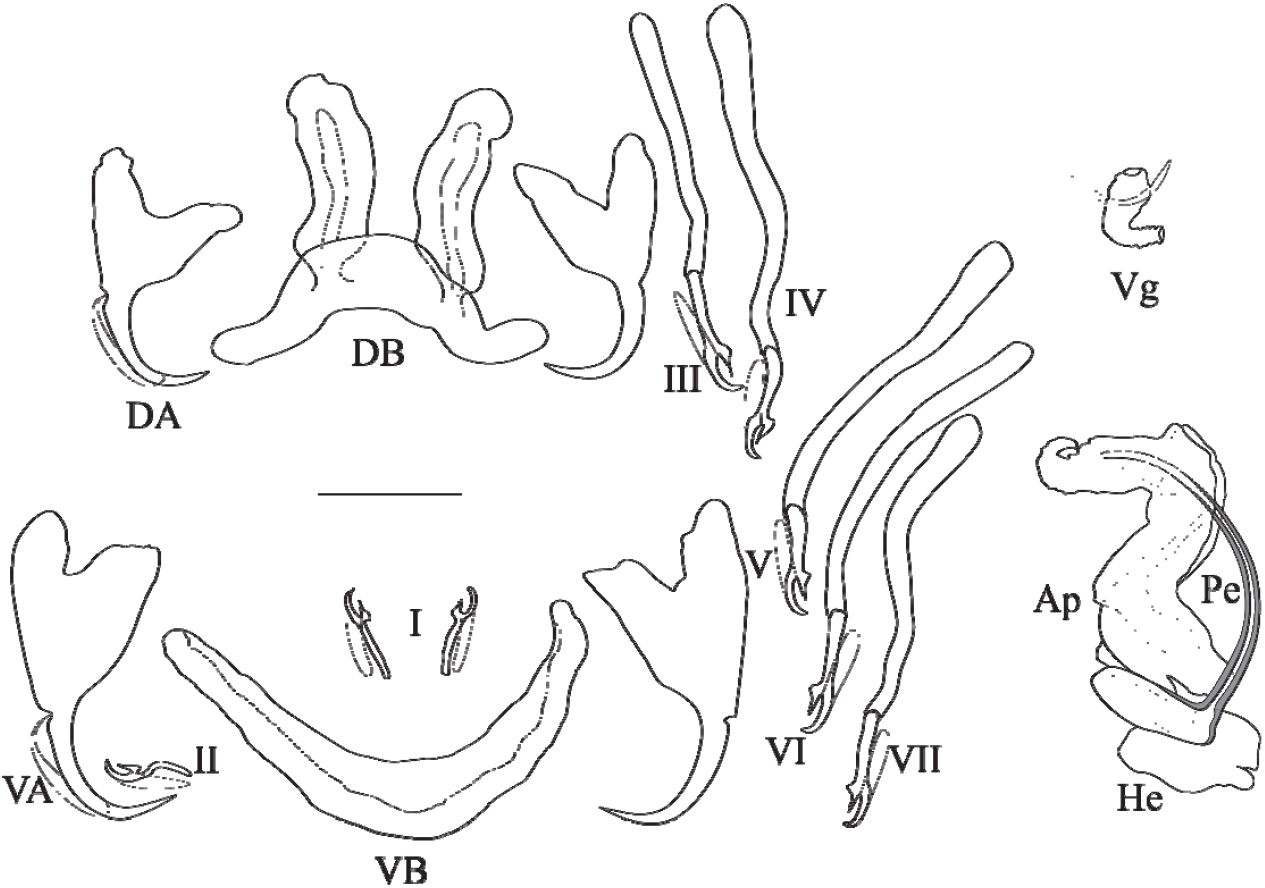
Sclerotized parts (haptor and male copulatory complex) of *Cichlidogyrus pseudodossoui* n. sp. **I–VII,** hook pairs; **DA,** dorsal anchors; **DB,** dorsal transverse bar; **VA,** ventral anchors; **VB,** ventral transverse bar. Male copulatory complex: **AP,** accessory piece; **Pe,** penis; **Vg,** vagina. Scale bar: 20 μm.

**Table 3.** Measurements of the new species of *Cichlidogyrus* described from haplochromine host species of Lake Victoria (N = number of specimens, n = number of observations per trait per species). All measurements (terminology following ICOPA IV, [1]) in μm, given as the mean and range (in parentheses)

| N | trait | *C. pseudodossoi* n. sp. |  | *C. nyanza* n. sp. |  | *C. cf. furu* n. sp. |  | *C. furu* n. sp. |  | *C. vetusmolendarius* n. sp. |  |
| --- | --- | --- | --- | --- | --- | --- | --- | --- | --- | --- | --- |
|  |  | 10<br>mean (min-max) | n | 54<br>mean (min-max) | n | 1<br>mean (min-max) | n | 36<br>mean (min-max) | n | 10<br>mean (min-max) | n |
| MCC | Ap | 36.2 (25.7-49.8) | 11 | 36.8 (22.2-46.7) | 48 | 36.5 (36.5-36.5) | 1 | 36.5 (15.4-45.3) | 34 | 19.6 (17.5-22.3) | 5 |
|  | He | 7.8 (6.7-8.8) | 3 | 1.5 (0.6-5.4) | 31 | 4.8 (4.8-4.8) | 1 | 5.6 (1.3-7.7) | 34 | 3.6 (3.0-5.0) | 7 |
|  | Pe | 58.5 (39.9-68.3) | 10 | 44.4 (35.7-66.4) | 48 | 58.6 (58.6-58.6) | 1 | 48.9 (23.7-66.1) | 34 | 27.8 (19.9-32.1) | 7 |
| DA | a | 29.7 (25.6-36.6) | 5 | 32.9 (29.3-35.9) | 37 | 37.6 (37.6-37.6) | 1 | 30.7 (18.5-35.1) | 29 | 38 (35.7-40.9) | 6 |
|  | b | 23.7 (22.9-24.3) | 5 | 25.7 (17.2-30.5) | 37 | 26.1 (26.1-26.1) | 1 | 21.5 (15.5-26.2) | 29 | 24.1 (20.0-26.2) | 6 |
|  | c | 11.4 (5.5-16.0) | 6 | 4.5 (1.9-10.8) | 37 | 4.8 (4.8-4.8) | 1 | 5.1 (1.8-12.5) | 29 | 6.1 (3.7-9.8) | 6 |
|  | d | 9.5 (1.6-13.1) | 6 | 10.5 (6.7-17.1) | 37 | 14.4 (14.4-14.4) | 1 | 11.9 (2.9-16.4) | 28 | 16.2 (13.5-18.5) | 6 |
|  | e | 6.4 (5.2-8.0) | 5 | 8.0 (5.7-9.6) | 36 | 7.0 (7.0-7.0) | 1 | 7.0 (4.3-9.4) | 28 | 7.7 (6.4-9.2) | 6 |
| DB | h | 23.2 (18.8-29.9) | 11 | 18.1 (12.6-23.6) | 72 | 16.7 (16.7-16.7) | 2 | 17.0 (10.9-23.2) | 54 | 15.7 (14.1-18.8) | 14 |
|  | w | 9.6 (5.9-28.6) | 11 | 7.9 (4.7-10.3) | 40 | 7.6 (7.6-7.6) | 1 | 7.1 (4.6-8.4) | 28 | 6.3 (4.0-8.3) | 8 |
|  | x | 47.0 (40.5-58.7) | 7 | 46.4 (27.4-57.5) | 42 | 32.4 (32.4-32.4) | 1 | 40.1 (33.0-49.3) | 29 | 40.4 (29.5-46.6) | 7 |
|  | y | 12.0 (9.0-13.9) | 9 | 13.9 (9.5-17.3) | 40 | 13.0 (13.0-13.0) | 1 | 12.4 (9.6-15.9) | 30 | 14.0 (13-15.9) | 7 |
| H | I | 10.9 (6.8-15.4) | 8 | 10.4 (7.0-13.9) | 59 | 11.8 (11.8-11.8) | 2 | 10.9 (9.3-12.2) | 45 | 27.2 (19.1-32.5) | 15 |
|  | II | 13.6 (11.9-14.9) | 0 | 13.3 (10.2-17.2) | 13 | 12.8 (12.8-12.8) | 1 | 15.8 (11.3-24.6) | 12 | 18.0 (18.0-18.0) | 1 |
|  | III-VII | 58.1 (48.3-66.4) | 37 | 16.8 (12.1-19.8) | 141 | 19.8 (19.8-19.8) | 4 | 18.1 (14.1-21.2) | 99 | 22.7 (18.9-26.2) | 29 |
| VA | a | 36.3 (33.0-41.0) | 7 | 33.3 (29.4-37.9) | 39 | 35.8 (35.8-35.8) | 1 | 31.7 (28.9-35.2) | 32 | 34.8 (31.4-39.9) | 9 |
|  | b | 30.1 (26.5-32.9) | 7 | 30.2 (26.6-33.8) | 38 | 31.9 (31.9-31.9) | 1 | 27.4 (21.9-31.4) | 30 | 29.4 (26.7-31.4) | 9 |
|  | c | 8.6 (6.6-13.0) | 7 | 4.0 (1.5-10.6) | 38 | 6.4 (6.4-6.4) | 1 | 4.3 (1.6-9.1) | 30 | 5.0 (2.3-6.4) | 9 |
|  | d | 12.1 (8.3-15.9) | 7 | 10.4 (4.4-15.2) | 40 | 11.7 (11.7-11.7) | 1 | 9.5 (3.3-13.6) | 30 | 9.5 (6.0-13.4) | 9 |
|  | e | 9.1 (7.5-11.4) | 7 | 10.7 (7.8-12.9) | 39 | 7.6 (7.6-7.6) | 1 | 8.9 (4.5-12.4) | 30 | 10.8 (9.2-12.6) | 9 |
| VB | w | 7.5 (6.0-8.4) | 6 | 8.9 (4.5-12.3) | 38 | 6.5 (6.5-6.5) | 1 | 6.2 (4.4-8.8) | 29 | 6.0 (5.1-7.0) | 8 |
|  | x | 41.8 (37.7-49.4) | 6 | 39.7 (27.4-46.6) | 40 | 40.7 (40.7-40.7) | 1 | 41.7 (30.6-51) | 29 | 47.4 (39.7-52.4) | 8 |
| Vg | L | 12.9 (11.0-16.2) | 8 |  |  |  |  |  |  |  |  |
|  | l | 5.3 (3.7-6.7) | 8 |  |  |  |  |  |  |  |  |

*Remarks:* this new species belongs to the group showing the following characters: hooks I short, long hooks III-VII, penis J-shaped, accessory piece Z-shaped without auxiliary plate, and sclerotised vagina. Previously described *C. thurstonae* and *C. tiberianus* from Lake Victoria (on few Oreochromini species and on a Haplochromini species) also fall into this group, but *C. pseudodossoui* n. sp. differs from them in the shape of the vagina (not bent in *C. thurstonae,* looped in *C. tiberianus).* This group further comprises: *C. anthemocolpos* Dossou, 1982 (on a few coptodonine species in West Africa), *C. bonhommei* Pariselle & Euzet, 1998 (on heterotilapiine species in West Africa), *C. bouvii* Pariselle & Euzet, 1997 (on an oreochromine species in West Africa), *C. dossoui* Douëllou 1993 (on several coptodonine species, a haplochromine, a pelmatolapiine, a tilapiine species in Central and West Africa, and introduced with oreochromines elsewhere), *C. douellouae* Pariselle, Bilong Bilong & Euzet, 2003 (on an oreochromine in West Africa), *C. ergensi* Dossou, 1982 (on several coptodonine species, a pelmatolapiine species in West Africa and Middle East), *C. flexicolpos* Pariselle & Euzet, 1995 (on several coptodonine species and a pelmatolapiine species in West Africa), *C. gillesi* Pariselle, Bitja Nyom & Bilong Bilong, 2013 (on a coptodonine species in Central Africa), *C. hemi* Pariselle & Euzet, 1998 (on a tilapiine species in West Africa), *C. kouassii* N’Douba, Thys van den Audenaerde & Pariselle, 1997 (on a coptodonine species in Southern-West Africa), *C. legendrei* Pariselle & Euzet, 2003 (on a pelmatolapiine species in Central Africa), *C. lemoallei* Pariselle & Euzet, 2003 (on few pelmatolapiini species in Central Africa), *C. ouedraogoi* Pariselle & Euzet, 1996 (on few coptodonine and a pelmatolapiine species in Central and West Africa), *C. testificatus* Dossou, 1982 (on a pelmatolapiine species in Central Africa), *C. tiberianus* Paperna, 1960 (on several coptodonine and Oreochromini species, few Haplochromini and tilapiine species, a pelmatolapiine species in Central and West Africa and in Middle East) and *C. vexus* Pariselle & Euzet, 1995 (on few coptodonine species in West Africa). In this morphological group, species differences can be seen in the shape of the vagina. *C. pseudodossoui* n. sp. mainly differs from *C. flexicolpos, C. lemoallei* and *C. testificatus,* by the shape and length of the vagina, very long and thin vs. short and bent at right angle. *C. pseudodossoui* n. sp. mainly differs from *C. ergensi, C. gillesi, C. kouassii* and *C. ouedraogoi* by the shape of the vagina, S-shaped vs. bent at right angle. *C. pseudodossoui* n. sp. mainly differs by the shape of the vagina (bent at right angle) from *C. anthemocolpos* (U-shaped with a distal plate), *C. bonhommei* (thin walled, bent in two different perpendicular plans), *C. hemi* (straight, with constant diameter and annulated all along), *C. legendrei* (well developed), *C. tiberianus* (looped, 1 turn). *C. pseudodossoui* n. sp. is close to *C. bouvii, C. dossoui, C. douellouae* and *C. vexus* which all have a conically shaped vagina bending at a right angle, but differs from these species by the greater length of its dorsal bar auricles and above all of its hook pairs III-VII (which have a size range of 58-66 in *C. pseudodossoui* n. sp. vs. 31-43, 36-50, 27-37, 31-43 respectively for the four other species). *Cichlidogyrus pseudodossoui* n. sp. was reported as *Cichlidogyrus* sp. III in [12–14]).

### *Cichlidogyrus nyanza* n. sp

urn:lsid:zoobank.org:act:3CCDC28A-959D-4336-B752-951B0F0DFCC7

*Type host: “Haplochromis” cyaneus* Seehausen, Bouton & Zwennes, 1998.

*Other hosts: Paralabidochromis chilotes* Greenwood 1959; *“Haplochromis” cyaneus* Seehausen, Bouton & Zwennes, 1998; *Ptyochromis xenognathus* Greenwood 1957; *P* sp. ‘striped rock sheller’; *Labrochromis* sp. ‘stone’; *Mbipia lutea* Seehausen & Bouton 1998; *M. mbipi* Seehausen, Lippitsch & Bouton 1998; *Neochromis gigas* Seehausen & Lippitsch 1998; *N. omnicaeruleus* Seehausen & Bouton, 1998; *N. rufocaudalis* Seehausen & Bouton 1998; *N.* sp. ‘unicuspid scraper’; *Pundamilia pundamilia* Seehausen & Bouton, 1998; *P.* sp. ‘pundamilia-like’; *P. nyererei* (Witte-Maas & Witte, 1985); *P.* sp. ‘nyererei-like’; *Pundamilia* sp. ‘Luanso’; *P.* sp. ‘pink anal’; *Pseudocrenilabrus multicolor victoriae* Seegers 1990.

*Infection parameters:* 4 of 4 *Paralabidochromis chilotes* from Makobe infected with 1 individual each, 5 of 5 *“Haplochromis” cyaneus* from Makobe infected with 1-5 individuals (mean intensity 3.0), 4 of 5 *Ptyochromis xenognathus* from Kissenda infected with 1-5 individuals (mean intensity 3.2), 1 of 1 *P.* sp. ‘striped rock sheller’ from Makobe infected with 3 individuals, 1 of 2 *Labrochromis* sp. ‘stone’ from Makobe infected with 3 individuals, 3 of 3 *Mbipia lutea* from Makobe infected with 3-5 individuals (mean intensity 4.0), 7 of 11 *M. mbipi* from Makobe infected with 1-3 individuals (mean intensity 1.7), 3 of 3 *Neochromis gigas* from Makobe infected with 4-6 individuals (mean intensity 5.0), 12 of 13 *N. omnicaeruleus* from Makobe infected with 1-17 individuals (mean intensity 4.1), 4 of 4 *N. rufocaudalis* from Makobe infected with 1-4 individuals (mean intensity 2.7), 19 of 22 *N.* sp. ‘unicuspid scraper’ from Makobe infected with 1-6 individuals (mean intensity 2.0), 13 of 14 *Pundamilia* sp. ‘Luanso’ from Luanso infected with 1-7 individuals (mean intensity 2.7), 5 of 24 *P. nyererei* from Makobe Island infected with 1-3 individuals (mean intensity 1.8), 9 of 16 *P.* sp. ‘nyererei-like’ from Kissenda infected with 1-2 individuals (mean intensity 1.4), 5 of 7 *P.* sp. ‘pink anal’ from Makobe infected with 1-5 individuals (mean intensity 1.8), 15 of 23 *P. pundamilia* from Makobe infected 1-6 with individuals (mean intensity 2.4), 9 of 14 *P.* sp. ‘pundamilia-like’ from Kissenda infected with 1-3 individuals (mean intensity 1.6), 1 of 1 *P.* sp. ‘pundamilia-like’ from Python infected with 3 individuals, 1 of 6 *Pseudocrenilabrus multicolor victoriae* from Sweya infected with 1 individual.

*Site:* Gills.

*Type locality:* Makobe Island (−2.3654, 32.9228).

*Other localities:* Kissenda Island (−2.5494, 32.8276), Python Island (−2.6237, 32.8567), Luanso Island (−2.6889, 32.8842).

*Material:* 54 whole-mounted specimens in Hoyer’s solution.

*Holotype:* MHNHxxxx-xx.

*Paratypes:* MHNHxxxx-xx. RMCA_VERMES_43420, SAMC-A092085.

*Symbiotype: “Haplochromis” cyaneus* Seehausen, Bouton & Zwennes, 1998 from Makobe (EAWAG ID 105317, **Table S1**).

*Symbioparatypes: “Haplochromis” cyaneus* Seehausen, Bouton & Zwennes, 1998 from Makobe (EAWAG ID 105323 and 105128, **Table S1**).

*Etymology:* The species epithet, a noun in apposition, refers to the word for large lake in several regional languages of East Africa.

*Description* (**Table 3**, **Fig. 7**): Two pairs of anchors of equal size and shape, with short shaft, dorsal anchor sometimes with fenestrations at the junction of shaft and guard. Ventral transverse bar U-shaped with thick branches. Dorsal transverse bar with two auricles inserted at its dorsal surface. Hooks 7 pairs; I and III to VII short (see [36, 37]). MCC consisting of a large penis with constant diameter, bent at distal third, accessory piece simple, S-shaped, attached to the basal bulb of the penis by a thin filament; no heel. No sclerotised vagina.

**Figure 6.7.**
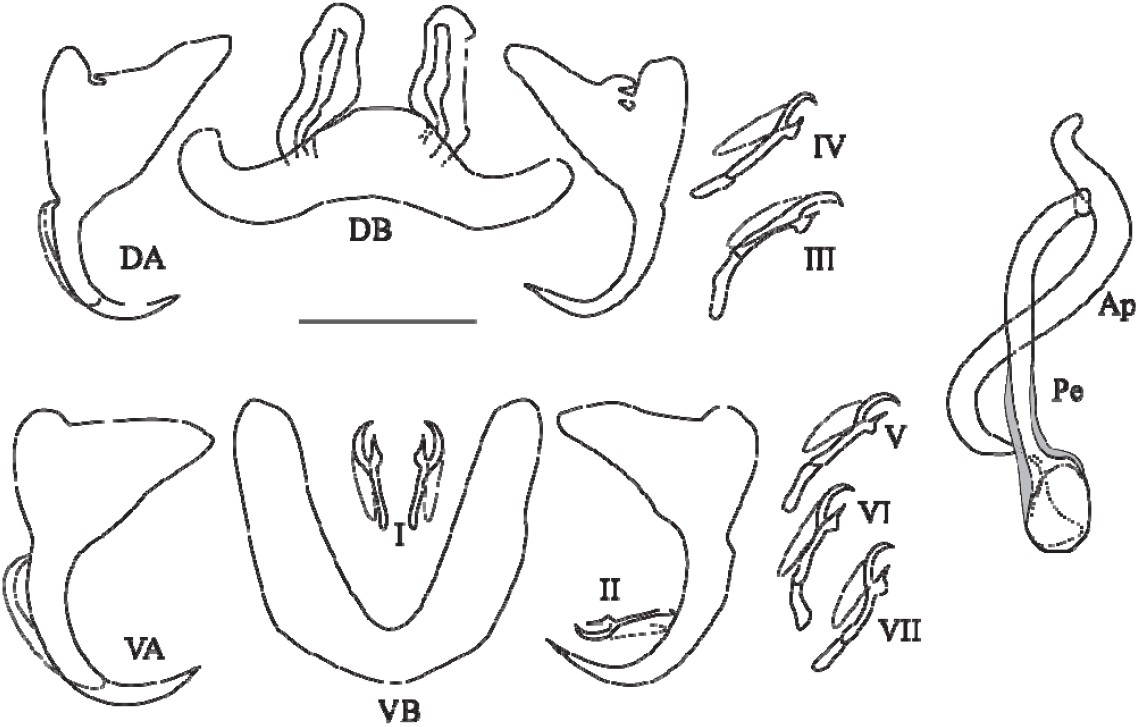
Sclerotized parts (haptor and male copulatory complex) of *Cichlidogyrus nyanza* n. sp. **I–VII,** hook pairs; **DA,** dorsal anchors; **DB,** dorsal transverse bar; **VA,** ventral anchors; **VB,** ventral transverse bar. Male copulatory complex: **AP,** accessory piece; **Pe,** penis. Scale bar: 20 μm.

*Remarks:* Short hook pairs I and III-VII, a simple MCC, and the absence of a sclerotized vagina are features that *Cichlidogyrus nyanza* n. sp. shares with several congeners infecting haplochromine cichlids in Central and Southern-East Africa, such as *C. gistelincki* Gillardin, Vanhove, Pariselle, Huyse & Volckaert, 2012; *C. irenae* Gillardin, Vanhove, Pariselle, Huyse & Volckaert, 2012; *C. steenbergei* Gillardin, Vanhove, Pariselle, Huyse & Volckaert, 2012; *C. banyankimbonai* Pariselle & Vanhove, 2015; *C. frankwillemsi* Pariselle & Vanhove, 2015; *C. franswittei* Pariselle & Vanhove, 2015; *C. muterezii* Pariselle & Vanhove 2015; *C. raeymaekersi* Pariselle & Vanhove, 2015; and *C. gillardinae* Muterezi Bukinga, Vanhove, Van Steenberge & Pariselle, 2012. However, this new species differs from all of these in showing the following combination of characters: hook I and III-VII short, accessory piece reaching beyond the distal end of the penis, no heel, no sclerotised vagina. Cichlidogyrus *haplochromii* and *C. tilapiae,* previously reported from Lake Victoria (on several haplochromine and few oreochromine species) also fall into the group with short hooks and simple male copulatory organ, but *C. nyanza* n. sp. differs from them in the shape of the accessory piece (of which the terminal end is blunter in *C. nyanza* n. sp. than the finger-like extension of *C. haplochromii* or the pointed extremity in *C. tilapiae)* and the shape of the ventral anchors (less incised in *C. nyanza* n. sp.). *Cichlidogyrus nyanza* n. sp. was reported as *Cichlidogyrus* sp. I in [12–14].

### *Cichlidogyrus furu* n. sp

urn:lsid:zoobank.org:act:C1E2FD2F-0BB1-4154-8C96-552C261283CC

*Type host: Pundamilia nyererei* (Witte-Maas & Witte, 1985).

*Other hosts: Astatoreochromis alluaudi* Pellegrin, 1904; *Paralabidochromis chilotes* Greenwood 1959; *“Haplochromis” cyaneus* Seehausen, Bouton & Zwennes, 1998; *Astatotilapia nubila* (Boulenger, 1906); *Ptyochromis xenognathus* Greenwood 1957; *Mbipia lutea* Seehausen & Bouton 1998; *M. mbipi* Seehausen, Lippitsch & Bouton 1998; *Neochromis omnicaeruleus* Seehausen & Bouton, 1998; *N. rufocaudalis* Seehausen & Bouton 1998; *N.* sp. ‘unicuspid scraper’; *Pundamilia* sp. ‘Luanso’; *Pundamilia nyererei* (Witte-Maas & Witte, 1985); *P.* sp. ‘pink anal’; *P. pundamilia* Seehausen & Bouton, 1998; *P.* sp. ‘pundamilia-like’ from Kissenda; *P.* sp. ‘nyererei-like’, *Pseudocrenilabrus multicolor victoriae* Seegers 1990.

*Infection parameters:* 2 of 9 *Astatoreochromis alluaudi* from Makobe infected with 1-2 individuals, 1 of 4 *Paralabidochromis chilotes* from Makobe infected with 1 individual, 1 of 5 *“Haplochromis “ cyaneus* from Makobe infected with 1 individual, 1 of 1 *Astatotilapia nubila* from Sweya infected with 3 individuals, 2 of 5 *Ptyochromis xenognathus* from Kissenda infected with 1-2 individuals (mean intensity 1.5), 1 of 3 *Mbipia lutea* from Makobe infected with 1 individual, 9 of 11 *M. mbipi* from Makobe infected with 1-3 individuals (mean intensity 1.8), 5 of 13 *Neochromis omnicaeruleus* from Makobe infected with 1-3 individuals (mean intensity 1.6), 2 of 4 *N. rufocaudalis* from Makobe infected with 1 individual each, 6 of 22 *N.* sp. ‘unicuspid scraper’ from Makobe infected with 1 individual each, 7 of 23 *Pundamilia pundamilia* from Makobe infected with 1-3 individuals (mean intensity 1.6), 6 of 14 *P.* sp. ‘pundamilia-like’ from Kissenda infected with 1-3 individuals (mean intensity 1.5), 15 of 24 *P. nyererei* from Makobe infected with 1-5 individuals (mean intensity 1.6), 7 of 14 *P.* sp. ‘nyererei-like’ from Kissenda infected with 1-3 individuals (mean intensity 1.6), 8 of 14 *P.* sp. ‘Luanso’ from Luanso infected with 1-5 individuals (mean intensity 2.1), 3 of 7 *P.* sp. ‘pink anal’ from Makobe infected with 1-5 individuals (mean intensity 2.6), 2 of 6 *Pseudocrenilabrus multicolor victoriae* from Sweya infected with 1 individual each.

*Site:* Gills.

*Type locality:* Makobe Island (−2.3654, 32.9228).

*Other localities:* Kissenda Island (−2.5494, 32.8276), Luanso Island (−2.6889, 32.8842), Sweya swampy inlet (−2.5841, 32.8970).

*Material:* 36 whole-mounted specimens in Hoyer’s solution.

*Holotype:* MHNHxxx.

*Paratypes:* MHNHxxx,.RMCA_VERMES_43418.

*Symbiotype: Pundamilia nyererei* (Witte-Maas & Witte, 1985) from Makobe (EAWAG ID 103397, **Table S1**).

*Symbioparatypes: Pundamilia nyererei* (Witte-Maas & Witte, 1985) from Makobe Island (EAWAG ID 103312 and 103397, **Table S1**).

*Etymology:* The species epithet is the word referring to haplochromine cichlids in Kiswahili, used as a noun in apposition.

*Description* (**Table 3**, **Fig. 8**): Two pairs of anchors of equal size and unequal shape (guard, and length difference between shaft and guard, more pronounced in dorsal anchors). Ventral transverse bar V-shaped with thin branches. Dorsal transverse bar thin with two auricles inserted at its dorsal surface. Hooks 7 pairs; I and III to VII short, except for pair V which is of medium size (see [36, 37]). Penis, S-shaped and sclerotized from its basal bulb until halfway its total length, is a tube which opens at its distal third and ends in a groove; accessory piece simple, attached to the rounded basal bulb of the penis by a thin filament; developed heel. No sclerotised vagina. Two specimens (deposited as vouchers under MHNHxxxx-xx and MHNHxxxx-xx), taken from one *P.* sp. ‘pink anal’ from Makobe (stored as 104372 at EAWAG) and one *P.* sp. ‘nyererei-like’ from Kissenda (stored as 104754 at EAWAG) differ from *C. furu* n. sp. in having the penis entirely sclerotized (**Fig. 9**, **Fig. 10**) but were otherwise very similar to *C. furu* n. sp..

**Figure 6.8.**
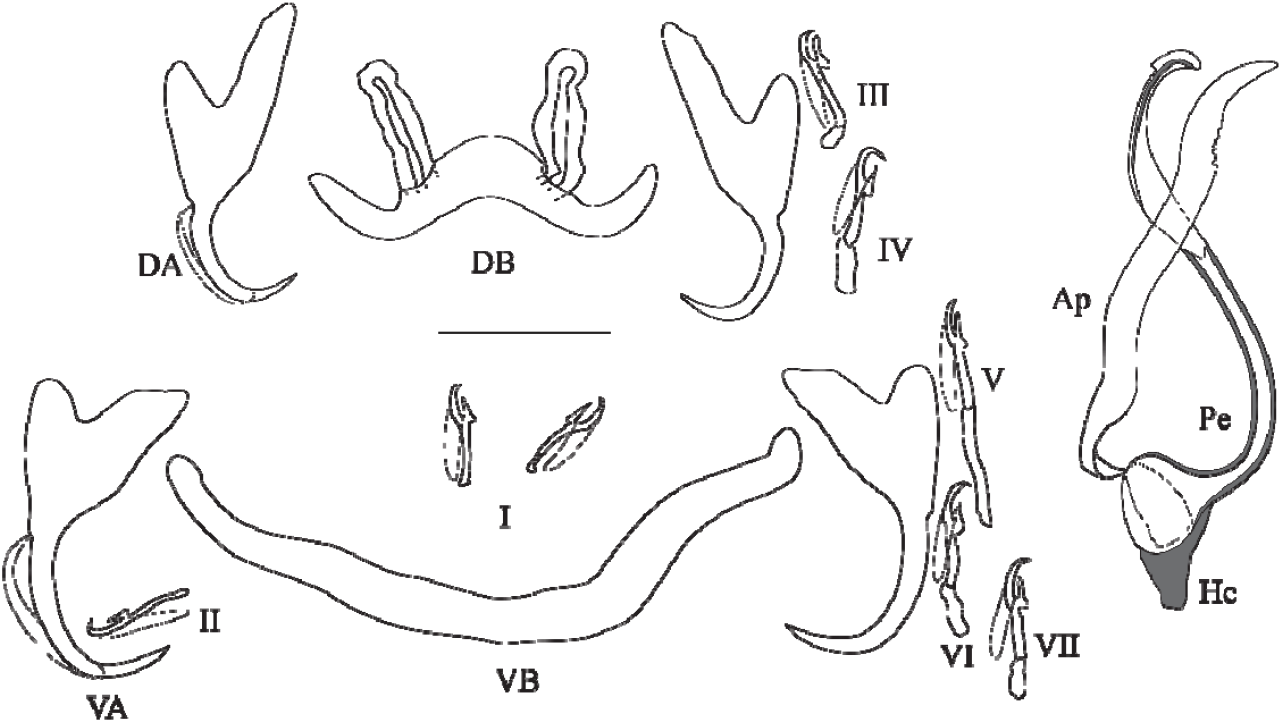
Sclerotized parts (haptor and male copulatory complex) of *Cichlidogyrus furu* n. sp. I–VII, hook pairs; **DA,** dorsal anchors; **DB,** dorsal transverse bar; **VA,** ventral anchors; **VB,** ventral transverse bar. Male copulatory complex: **AP,** accessory piece; **Pe,** penis. Scale bar: 20 μm.

**Figure 6.9.**
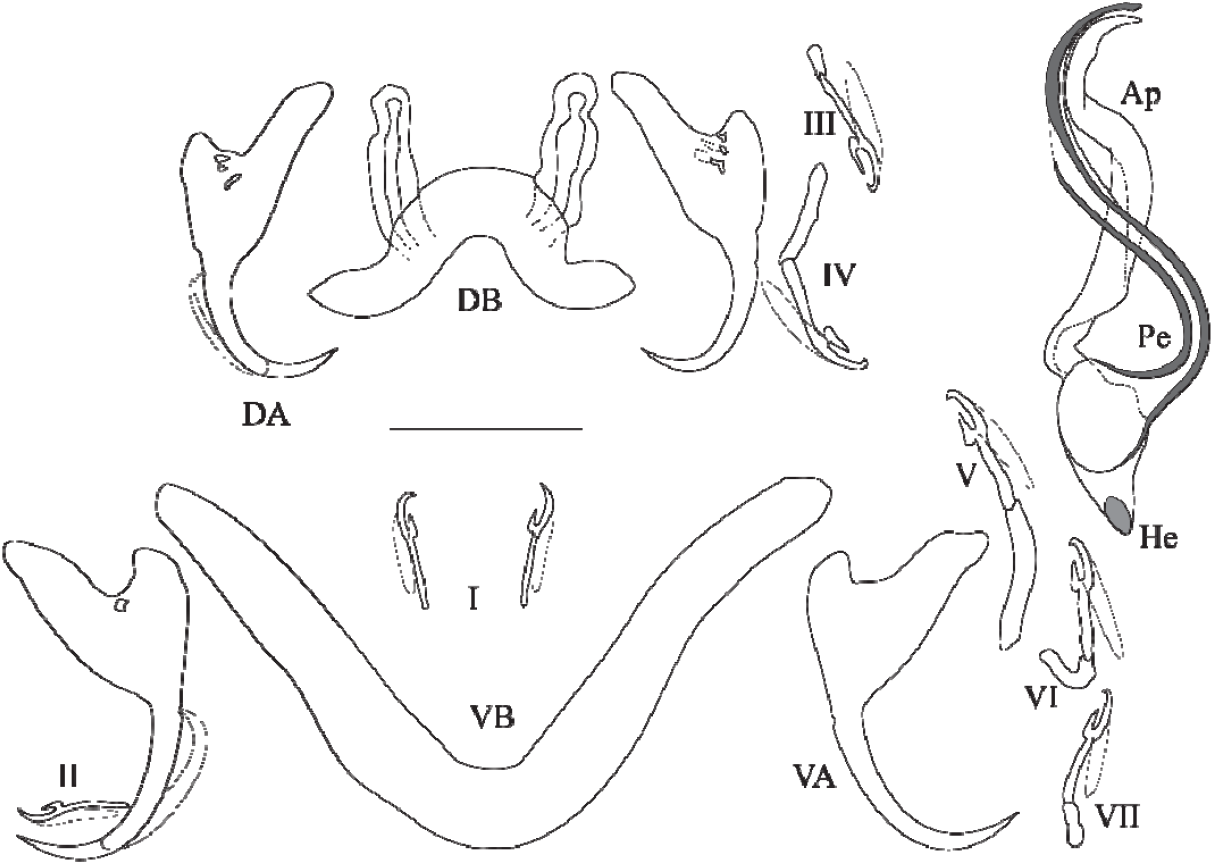
Sclerotized parts (haptor and male copulatory complex) of *Cichlidogyrus* cf. *furu* n. sp. **I–VII,** hook pairs; **DA,** dorsal anchors; **DB,** dorsal transverse bar; **VA,** ventral anchors; **VB,** ventral transverse bar. Male copulatory complex: **AP,** accessory piece; **Pe,** penis. Scale bar: 20 μm.

**Figure 6.10.**
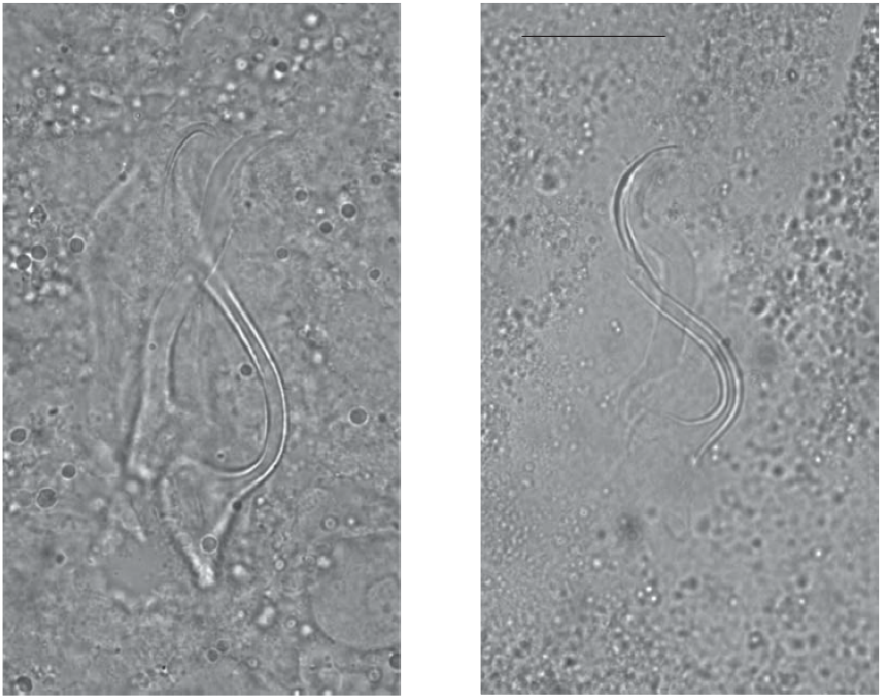
Micrographs of the male copulatory organ of *Cichlidogyrus furu* n. sp. **(left)** and of *C.* cf. *furu* n. sp. **(right),** fixed in Hoyer’s medium. Scale bar: 20 μm.

Since measurements of this morphotype fall into the range of *C. furu* n. sp. and we cannot exclude that the peculiar trait resulted from mounting, we report it as *Cichlidogyrus* cf. *furu* n. sp. and not as a new species.

*Remarks: C. furu* n. sp. is unique among species of *Cichlidogyrus* in having a penis ending in a groove. This feature had already been described in other dactylogyridean monogenean species (e.g. *Synodontella speroadotevii* Bouah, N’Douba & Pariselle 2019; [1]). *Cichlidogyrus furu* n. sp. was reported as *Cichlidogyrus* sp. II in [12–14].

### *Cichlidogyrus vetusmolendarius* n. sp

urn:lsid:zoobank.org:act:A526328E-270B-4B95-921C-FB1A3C528586

*Type host: Pseudocrenilabrus multicolor victoriae* Seegers 1990.

*Other hosts:* one *Astatotilapia nubila* (Boulenger, 1906); *Mbipia lutea* Seehausen & Bouton 1998; *Pundamilia pundamilia* Seehausen & Bouton, 1998; *P.* sp. ‘pundamilia-like’; *P. nyererei* (Witte-Maas & Witte, 1985); *P.* sp. ‘nyererei-like’; *P.* sp. ‘pink anal’.

*Infection parameters:* 1 of 1 *Astatotilapia nubila* from Sweya infected with 2 individuals, 1 of 3 *Mbipia lutea* from Makobe infected with 1 individual, 2 of 23 *Pundamilia pundamilia* from Makobe infected with 1 individual each, 1 of 15 *P.* sp. ‘pundamilia-like’ from Kissenda infected with 1 individual, 3 of 24 *P. nyererei* from Makobe infected with 1 individual each, 1 of 16 *P.* sp. ‘nyererei-like’ from Kissenda infected with 2 individuals, 1 of 7 *P.* sp. ‘pink anal’ from Makobe infected with 1 individual, 3 of 6 *Pseudocrenilabrus multicolor victoriae* from Sweya infected with 1 individual each.

*Site:* Gills.

*Type locality:* Makobe Island (−2.3654, 32.9228).

*Other localities:* Kissenda Island (−2.5494, 32.8276), Luanso Island (−2.6889, 32.8842), Sweya swampy inlet (−2.5841, 32.8970).

*Material:* 10 whole-mounted specimens in Hoyer’s solution.

*Holotype:* MHNHxxx.

*Paratypes:* MHNHxxxx, RMCA_VERMES_43422.

*Symbiotype: Pseudocrenilabrus multicolor victoriae* Seegers 1990 (EAWAG ID 106984, **Table S1**).

*Symbioparatypes: Mbipia lutea* Seehausen & Bouton 1998 from Makobe (EAWAG ID 13313, **Table S1**); *Astatotilapia nubila* (Boulenger, 1906) from Sweya (EAWAG ID 109405, **Table S1**).

*Etymology:* The species epithet refers to the type host, a representative of a relatively old haplochromine lineage compared to most other hosts studied (“vetus”, Latin adjective meaning “old”), and to the shape of the first pair of hooks, similar to the wings of a windmill (“molendarius”, Latin adjective derived from “molendinum”, “windmill”).

*Description* (**Table 3, Fig. 11**): Two pairs of anchors of unequal shape and size (guard longer in dorsal anchors). Ventral transverse bar long and thin, V-shaped. Dorsal transverse bar thin with two auricles inserted at its dorsal surface. Hooks 7 pairs; I thick and long; III to IV short, V to VII of medium size (see [36, 37]). Penis short, thin, straight and spirally coiled (1.5 turns); accessory piece simple, attached to the basal bulb of the penis by a thin filament, spirally coiled (1.5 turns) winding around the penis; poorly developed heel. No sclerotised vagina.

**Figure 6.11.**
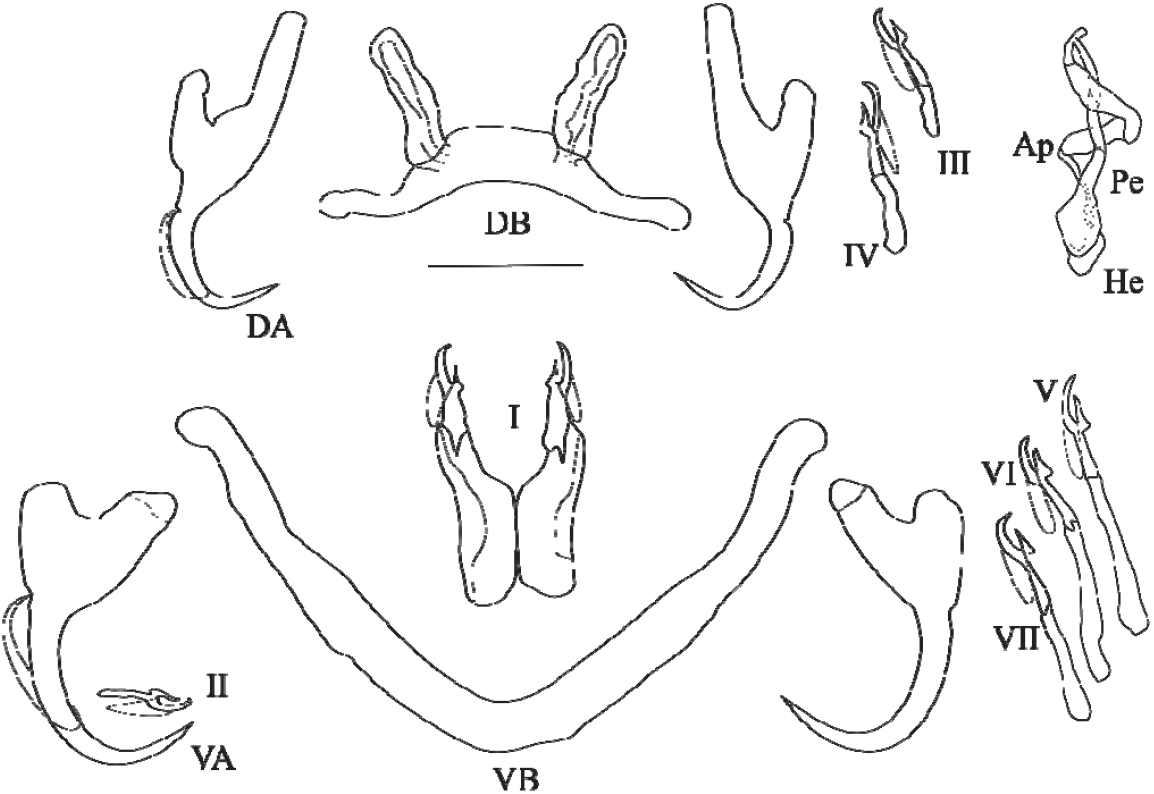
Sclerotized parts (haptor and male copulatory complex) of *Cichlidogyrus vetusmolendarius* n. sp. **I–VII,** hook pairs; **DA,** dorsal anchors; **DB,** dorsal transverse bar; **VA,** ventral anchors; **VB,** ventral transverse bar. Male copulatory complex: **AP,** accessory piece; **Pe,** penis. Scale bar: 20 μm.

*Remarks:* this new species belongs to the group showing the following characters: hooks pair I very large, no visible vagina. The previously described *C. dionchus* Paperna & Thurston, 1969 from Lake Victoria haplochromines falls into this morphological group, but it differs from *C. vetusmolendarius* n. sp. in the shape of its accessory piece (which is curved instead of coiled in *C. vetusmolendarius* n. sp., and broadens terminally ending in a hook-like extremity), its clearly elongated heel (poorly developed in *C. vetusmolendarius* n. sp.) and its dorsal anchors with proportionally longer point and blade. This group also comprises: *C. berradae* Pariselle & Euzet, 2003 (on a few coptodonine and a pelmatolapiine species of Central Africa), *C. bulbophallus* Geraerts & Muterezi Bukinga, 2020 (on a haplochromine species in Central Africa), *C. digitatus* Dossou, 1982 (on several coptodonine, a pelmatolapiine and a Tilapiine species in Central and West Africa), *C. halinus* Paperna, 1969 (on an oreochromine species in West Africa), *C. maeander* Geraerts & Muterezi Bukinga, 2020 (on a haplochromine and a tilapiine species in Central Africa), *C. nuniezi* Pariselle & Euzet, 1998 (on a tilapiine species in West Africa), *C. papernastrema* Price, Peebles & Bamford, 1969 (on a coptodonine, a tilapiini and an oreochromini species in Central and Southern Africa), *C. philander* Douëllou, 1993 (on a haplochromine in Southern Africa), *C. quaestio* Douëllou, 1993 (on a few haplochromine, a tilapine and a coptodonine species in Southern Africa) and *C. yanni* Pariselle & Euzet, 1995 (on several coptodonine species and a pelmatolapiine species in West Africa). Among these species, only *C. maeander* has a spirally coiled accessory piece winding around the penis like *C. vetusmolendarius* n. sp.. Both species share features of the haptoral sclerotised parts (guards about twice as long as shafts in both anchors, with the guard more pronounced in the dorsal anchors; dorsal anchors on average longer than ventral ones; auricles implanted at the dorsal surface of the dorsal bar; long first hook pair), and their accessory pieces resemble each other. Some measurements distinguish *C. vetusmolendarius* n. sp. from *C. maeander:* ventral anchor length (33-36 vs 43-44 μm), hook pair I length (25-32 vs. 32-41). The main difference concerns the shape of the penis: “stylet short, forming enlarged bulb at base; base attached to pronounced heel; penis stylet distally curved, with pointed end” for *C. maeander* (Geraerts et al., 2020) vs. “short, thin, straight and spirally coiled (1.5 turns); “poorly developed heel” for *C. vetusmolendarius* n. sp.. *Cichlidogyrus vetusmolendarius* n. sp. was reported as *Cichlidogyrus* sp. V in [12–14].

## Discussion

Recent research on the evolutionary ecology of cichlid-parasite interactions in Lake Victoria proposed that the communities of dactylogyrid flatworms infecting the lake’s haplochromine radiation differ from those parasitizing species of two ancient non-radiating haplochromine lineages in the same locations [12]. Cichlid species that are members of the young radiation seemed to share the same set of dactylogyrid flatworm species among each other but also to some extent with the ancient haplochromine lineages. Here we aimed to underpin these observations with formal taxonomic assessment of the gill monogeneans in question. Of the ten species previously reported in Lake Victoria, we only found two in the present study *(Cichlidogyrus bifurcatus* and *C. longipenis).* This can be explained by the difference in locations and/or host lineages surveyed. We surveyed Haplochromini from the southern part of Lake Victoria (Tanzania), while previous studies surveyed the northern part (Uganda) and mostly focused on other tribes (such as Oreochromini and Tilapiini) [33, 35, 34]. We may speculate that host lineage, rather than geographic locality, is the more important factor in determining infection variation because the two abovementioned species of *Cichlidogyrus* have been documented in the same host species in the present study and in the previous ones. This study increases the number of nominal monogenean species that are exclusively known from Lake Victoria from one (*C. longipenis)* to five. The fact that only four (or five, when including *C.* cf. *furu* n. sp.) new species are found when characterising the monogenean fauna of 20 host species, from five different locations, is in line with the trends suggested by Pariselle et al. [39]: the discovery rate and proportion of endemism in *Cichlidogyrus* are lower in the young Lake Victoria compared to the ancient Lake Tanganyika. Indeed, when comparing Lake Victoria’s littoral haplochromines with Tropheini, a lineage of littoral Tanganyika haplochromines, the species richness and uniqueness of their monogenean parasites are starkly lower in the former (see e.g. [57], indicating typically at least one, and up to seven, unique parasite species per host species, with only closely related congeneric hosts, rarely, sharing parasite species). This contrasts with the species richness of the cichlid hosts which is twice higher in Lake Victoria than in Lake Tanganyika [54, 9]. Unlike the haplochromine cichlids in Lake Tanganyika, Lake Victoria haplochromines were now found to also harbour species displaying some long hook pairs and a more complex accessory piece of the MCC (*C. pseudodossoui* n. sp. and *C. vetusmolendarius* n. sp.). Furthermore, it is the first time that a penis ending in an open groove was observed in *Cichlidogyrus,* in *C. furu* n. sp., adding to the diversity in the morphology of monogeneans infecting African Great Lake haplochromines. Also noteworthy about *Cichlidogyrus furu* n. sp. is that, while the relative lengths of hook pairs III to VII were previously often considered together [36, 59], in this species, pair V is clearly longer than the others. Size differences within hook pairs III to VII have also been reported from recently discovered congeners in the Congo river basin [18, 10]. In agreement with Rahmouni et al. [42] and Geraerts et al. [10], this further illustrates that the different haptor configurations proposed for the classification of species of *Cichlidogyrus* into four morphotype groups, mostly based on West African representatives (see [40, 59]) cannot accommodate all the recently described species from Central, East and Southern Africa. Phylogenetic analyses with increased species coverage are needed to re-evaluate to what extent the relative length of the haptoral hooks is systematically informative. The genetic data required for this could also help in clarifying whether some morphological features could be key to understand the shared history of these species. This may be the case for the tube-shaped penis and simple accessory piece of *C. bifurcatus, C. longipenis* and *C. nyanza* n. sp., that are shared with many congeners infecting haplochromines (see [11, 31, 55]), and for the long hooks V shared by *C. furu* n. sp., *C. calycinus,* and *C. omari* [18]. Pending confirmation of this phylogenetic signal, we note the morphological similarity of these species with typical parasites of haplochromines (reported from representatives of e.g. *Haplochromis* Hilgendorf, 1888, *Orthochromis* Greenwood, 1954, *Pharyngochromis* Greenwood, 1979, *Pseudocrenilabrus* Fowler, 1934, *Sargochromis* Regan, 1920, *Serranochromis* Regan, 1920 and Tropheini), namely a simple male copulatory organ and similar size and shape of ventral and dorsal anchors. On the other hand, *C. pseudodossoui* n. sp. resembles species described from mainly West African cichlid hosts belonging to Heterotilapiini, Pelmatolapiini, Gobiocichlini, Coptodonini and Oreochromini, and hence probably belongs to an entirely different lineage of *Cichlidogyrus.* This suggests that members of *Cichlidogyrus* colonised Lake Victoria haplochromines or their ancestors at least twice. The Lake Victoria region was colonised by two cichlid lineages, the upper Nile and the Congolese lineages, whereas Lake Victoria itself was colonised by at least four lineages belonging to two tribes (Haplochromini and Oreochromini) [27]. The taxonomic coverage of the current *Cichlidogyrus* phylogenetic reconstruction across Africa does not suffice to test whether this provides a likely pathway for the origins of the Lake Victoria cichlid monogeneans. However, in the context of the phylogeny of *Cichlidogyrus,* we can already observe that the haptors of the Lake Victoria parasites have auricles attached to the dorsal side of the dorsal bar. This is considered a derived state, in contrast to the auricles being a continuation of the anterior side of the dorsal bar, as in the earlier diverged parasites of the only very distantly related tylochromine cichlids [30]. While the dominance of *C. bifurcatus* on *P. multicolor victoriae,* and of *C. longipenis* on *A. alluaudi,* in contrast to the communities of *Cichlidogyrus* of the other investigated hosts clearly demonstrate community-level differences between member species of the haplochromine radiation and other Lake Victoria haplochromines, none of these monogeneans seems specialised to a single host species. We concur with Pariselle et al. [39] that this low host-specificity may be a consequence of the young age of the lake and its fish assemblage. It would therefore be interesting to expand sampling to adjacent lakes and river systems, to test the hypothesis that their haplochromine cichlids harbour the same or closely related species of *Cichlidogyrus* [62, 27].

## Acknowledgements

This research was funded by the University of Bern, the Swiss National Science Foundation and the University of Groningen (Ubbo Emmius Programme). Infrastructure was provided by the Natural History Museum in Lugano and Hasselt University (EMBRC Belgium - FWO project GOH3817N). MPMV received support from the Special Research Fund of Hasselt University (BOF20TT06). Sampling was conducted with permission of the Tanzania Commission for Science and Technology (COSTECH - No. 2013-253-NA-2014-117). We thank Chahrazed Rahmouni and Michiel Jorissen for insights on morphological identification.

